# Single-Cell Analysis of Gene Regulatory Networks in the Mammary Glands of P4HA1-knockout mice

**DOI:** 10.1101/2024.11.05.622022

**Authors:** Akshat Gupta, Lilin Huang, Jinpeng Liu, Ren Xu, Wei Wu

## Abstract

Prolyl hydroxylation, catalyzed by collagen prolyl-4 hydroxylase (P4H), is a crucial post-translational modification involved in collagen biosynthesis. P4HA1, an isoform of P4H, plays a prominent role in stabilizing hypoxia-inducible factor-1α (HIF-1α). P4HA1 is frequently upregulated in highly aggressive triple-negative breast cancer and has been implicated in tumor progression, metastasis, and chemoresistance.

In this study, we investigated the role of P4HA1 in mouse mammary glands by analyzing gene regulatory networks (GRNs) in basal epithelial cells across two mouse groups: control (5Ht) and P4HA1-knockout (6Ho) mice. Specifically, we employed a single-cell network inference approach, integrating single-cell RNA sequencing with the SCENIC pipeline, and incorporated multiple validation strategies to construct gene regulatory networks (GRNs) specific to basal epithelial cells from each mouse group. Despite the inherent challenges of single-cell data, our approach identified robust and reproducible GRN patterns across both mouse groups. Based on these patterns, we identified subclusters of basal epithelial cells with similar regulatory profiles across the two mouse groups and a unique subcluster in the control mice with a distinct regulatory pattern absent in the P4HA1-deficient 6Ho mice. This unique subcluster exhibited concurrent activation and potential crosstalks between stem cell development and inflammatory response pathways, underscoring the crucial role of P4HA1 in regulating these biological processes linked to cancer progression. We validated these findings through multiple approaches, including experimental validation. Given that the loss of P4HA1 disrupts the interplay between stem cells and inflammation, our results suggest that targeting P4HA1 may offer a promising therapeutic strategy for breast cancer treatment.

**Author Summary:** In this study, we aimed to understand how the P4HA1 gene affects the development and functions of mammary gland cells in mice using a single-cell network approach. P4HA1 plays an important role in collagen production, which is essential for maintaining the structure of normal tissues; however, it also helps stabilize the hypoxia-inducible factor-1α (HIF-1α) protein that responds to low oxygen levels, a condition often found in various cancers. We generated and analyzed single-cell RNA sequencing data to understand the regulatory roles of P4HA1 in two groups of mice: one with a single functional allele of P4HA1 and the other with the P4HA1 gene knocked out. We constructed gene regulatory networks for basal epithelial cells in both groups using the SCENIC pipeline. Due to the inherent complexity and noise within single-cell data, we employed multiple validation strategies to ensure that the identified regulatory patterns in the networks across both mouse groups were robust and reproducible.

In particular, we found that the P4HA1 gene plays a crucial role in pathways related to stem cell development and inflammation, which are essential for both tissue growth and repair, as well as cancer progression. One unique subcluster of basal epithelial cells found only in control mice with P4HA1 showed simultaneous activation of stem cell and inflammatory processes, suggesting that this gene plays a significant role in regulating the communication between these vital cellular processes. Our work provides new insights into how P4HA1 might serve as a therapeutic target to slow down or prevent the progression of aggressive cancers, such as triple-negative breast cancer, where treatment options are currently limited, leading to a high mortality rate.

## Introduction

Prolyl hydroxylation is a critical post-translational modification that enhances protein folding and stability, primarily by influencing collagen biosynthesis (1). This modification is catalyzed by the enzyme collagen prolyl-4 hydroxylase (P4H). We and others have shown that P4HA1, an isoform of P4H, enhances the stability of hypoxia-inducible factor-1α (HIF-1α), a vital regulator of the cellular response to hypoxia, which promotes angiogenesis, cancer metastasis, and chemoresistance (2, 3). P4HA1 is frequently upregulated in cancer, particularly in triple-negative breast cancer (TNBC), a subtype associated with poor prognosis and aggressive growth (4, 5).

In this study, we investigated the role of P4HA1 in mouse mammary glands by analyzing gene regulatory networks (GRNs) in basal epithelial cells across two distinct mouse groups. The first group, referred to as 5Ht, has one functional allele of P4HA1 and serves as the control, while the second group, referred to as 6Ho, has a P4HA1-knockout where P4HA1 expression is silenced in basal mammary epithelial cells.

The generation and inference of GRNs with single-cell transcriptomic data have long been challenging in computational biology. As the volume of RNA sequencing data grows, effective network reconstruction becomes increasingly important. GRNs serve as graphical representations of biological systems, where nodes represent genes, and edges denote regulatory interactions between these genes (6), providing insights into complex regulatory mechanisms underlying cellular behaviors.

The advent and widespread adoption of single-cell RNA sequencing (scRNA-seq) datasets have spurred the development of various network inference methods (7). The SILGGM approach leverages Gaussian graphical models (GGMs), which estimate conditional dependencies between genes by transforming the problem of network estimation into a sparse estimation of precision matrices (8). The information-incorporated Gaussian graphical model approach enhances traditional GGMs by integrating prior knowledge from existing studies, using penalization to balance observed data with supplementary information. This approach adaptively includes reliable gene connections while down-weighting potentially inaccurate information, improving inference accuracy and robustness (9). Information-theoretic approaches, on the other hand, use measures such as mutual information to capture both linear and non-linear relationships between genes (10). By considering higher-order interactions and multivariate dependencies, methods such as Partial Information Decomposition (PIDC) enable the inference of more complex regulatory networks by identifying synergistic and redundant information across groups of genes (10, 11). The Boolean networks method models genes as binary variables and their interactions are modeled using logical operators; this method captures the dynamic behavior of GRNs by simulating state transitions through asynchronous updates (12).

In several benchmark studies (7, 13), SCENIC (14) has emerged as a robust and scalable approach for gene network inference with single-cell data. SCENIC effectively handles large datasets, mitigates the effects of technical dropout standards in single-cell protocols, and produces reliable results without assuming specific data distributions. Specifically, SCENIC first utilizes non-linear, nonparametric models such as Gradient Boosting regression to infer co-expression-based networks. It then uses motif enrichment analysis to identify transcription factors (TFs) and their direct target genes, refining the co-expression networks into regulons (a collective term referring to TFs and their target genes). Lastly, the activities of these regulons in individual cells are measured using an Area Under the Curve (AUC) metric. This metric ranks all genes in a cell based on their expression levels and examines if the genes in a regulon are near the top of that ranking. The more highly ranked the genes in the regulon are, the more active the regulon is considered in that cell.

In this study, we generated gene regulatory networks specific to basal epithelial cells in the mammary glands of both the 5Ht and 6Ho mice using SCENIC, aiming to investigate the molecular mechanisms in these two groups. Despite the superior performance of SCENIC, the inherent limitations of single-cell data, such as low RNA content, frequent dropout events, and technical noise, pose challenges in accurately reconstructing reliable gene networks. These limitations complicate downstream analyses, particularly when comparing gene network results from the control and knockout mice. Disentangling actual biological variations from artifacts introduced by data quality was a major challenge that we encountered in this study.

To address these limitations, we utilized various validation strategies, including experimental validation, to enhance the reliability of our networks and findings. By combining single-cell network inference techniques with these validation approaches, we provide novel insights into the role of P4HA1 in regulating both stem cell differentiation and inflammation within basal epithelial cells of the mouse mammary gland. Our findings reveal a distinct regulatory subcluster in the basal epithelial cells of the 5Ht mice, where concurrent activation of stem cell development and inflammatory response pathways forms a mechanism potentially driving cancer progression (15–17). The absence of this subcluster in the P4HA1-deficient 6Ho mice suggests that removing P4HA1 disrupts these pathways in the knockout model. Given the known roles of inflammation and stem cell development in cancer progression (18), our results indicate that inhibiting P4HA1 could offer a promising therapeutic strategy for treating breast cancer.

## Results

In this study, we generated scRNA-seq data from the mammary glands of two groups of mice: the control (5Ht) mice and the P4HA1-knockout (6Ho) mice. We integrated their single-cell data using Seurat (19) to compare gene expression differences, shared cell states, and types between the two mouse groups. We also annotated basal and luminal epithelial cells and macrophages based on well-known cell markers (see Materials and Methods for details). The numbers of the cells in these three major cell types across both mouse groups are summarized in S1 Table.

The Uniform Manifold Approximation and Projection (UMAP) plot (20) in Fig 1A shows that scRNA-seq data from both groups integrate very well, overlapping nearly perfectly. Also, the three major cell types – basal and luminal epithelial cells and macrophages – segregate into distinct clusters (Fig 1B), consistent with previous studies (21, 22). To visualize and compare each mouse group individually while preserving the overall context of integration, we split the integrated UMAP plot by the mouse group. S1B Fig shows that the cell clusters in both the 5Ht and 6Ho mice remain highly similar in terms of shape and location, as expected from the well-overlapping integrated plot.

**Fig 1:**
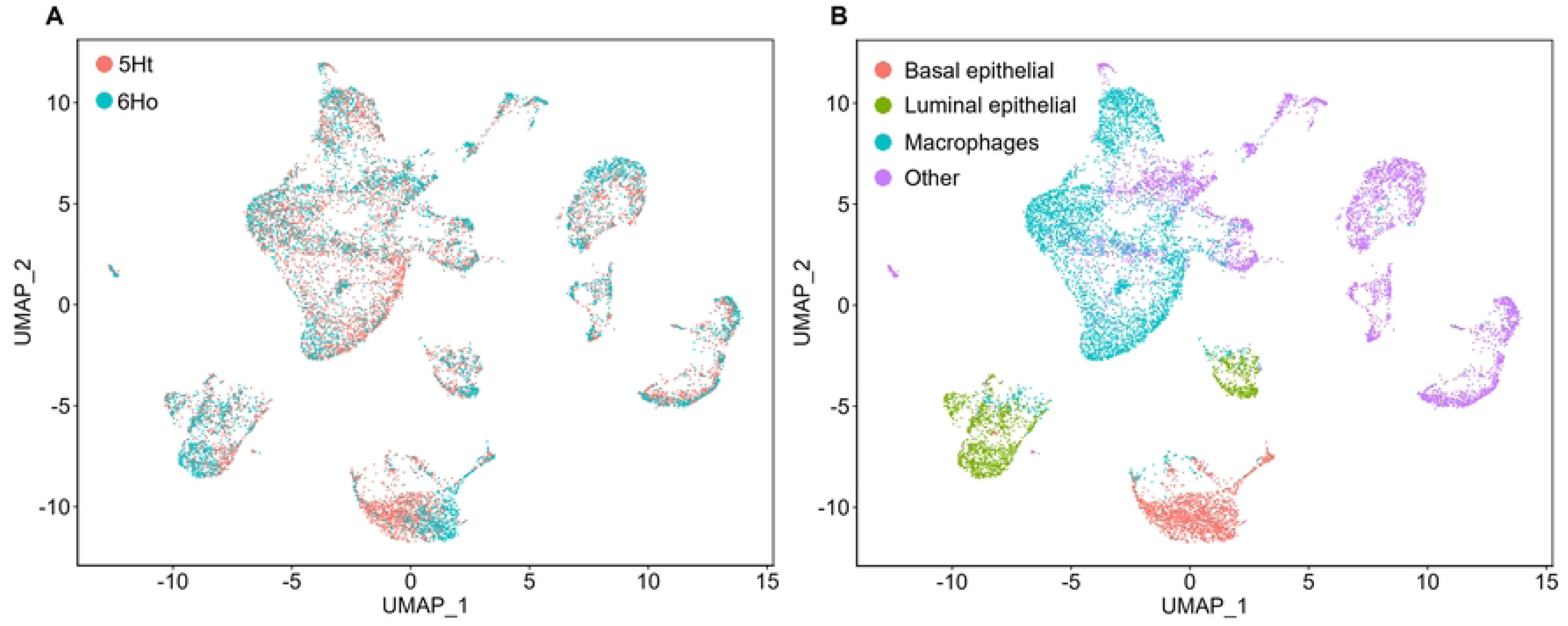
Integrated UMAP plots annotated by the 5Ht and 6Ho mouse groups (A) and by cell types (B).

### Network Analysis

Next, through network and cluster analyses, we aimed to identify the mechanistic differences underlying the mammary basal epithelial cells between the 5Ht and 6Ho mice. Using SCENIC, we identified 268 and 270 regulons activated in the 5Ht and 6Ho basal epithelial cells, respectively. Of these regulons, 222 were regulated by the common TFs between the two mouse groups (shown in S2 Table). The remaining regulons in the basal epithelial cells of each mouse group were identified as unique.

### Cluster Analysis

As shown by previous studies (22), mouse mammary basal epithelial cells can be characterized into several subclusters, which were identified using cluster analysis based on the gene expression levels derived from RNA-seq counts. Since our goal was to understand the mechanistic differences between the two mouse groups, we instead sought to identify subclusters of the basal epithelial cells based on the regulon activities in these mice. In particular, we utilized the regulons regulated by the 222 common TFs to perform cluster analysis; since these TFs were identified through network analysis in both the 5Ht and 6Ho mice, we considered their regulons more robust and reliable than the unique regulons detected in each mouse group.

#### Aligning basal epithelial subclusters in the 5Ht and 6Ho mice

Our cluster analysis revealed 5 basal epithelial subclusters in each mouse group (Fig 2). To compare them, we first needed to align the subclusters. While most of the cells in both mouse groups should be of the same types, the differences between the mouse groups may lead to distinct regulatory behaviors and gene expression patterns, making alignment challenging. As such, we reasoned that since basal epithelial cells in the 6Ho mice are P4HA1-deficient, some subclusters of cells should exhibit different regulatory patterns compared to their counterparts in the 5Ht mice, while others should remain similar. Thus, we employed the following strategy to align the basal epithelial subclusters in the two mouse groups.

**Fig 2:**
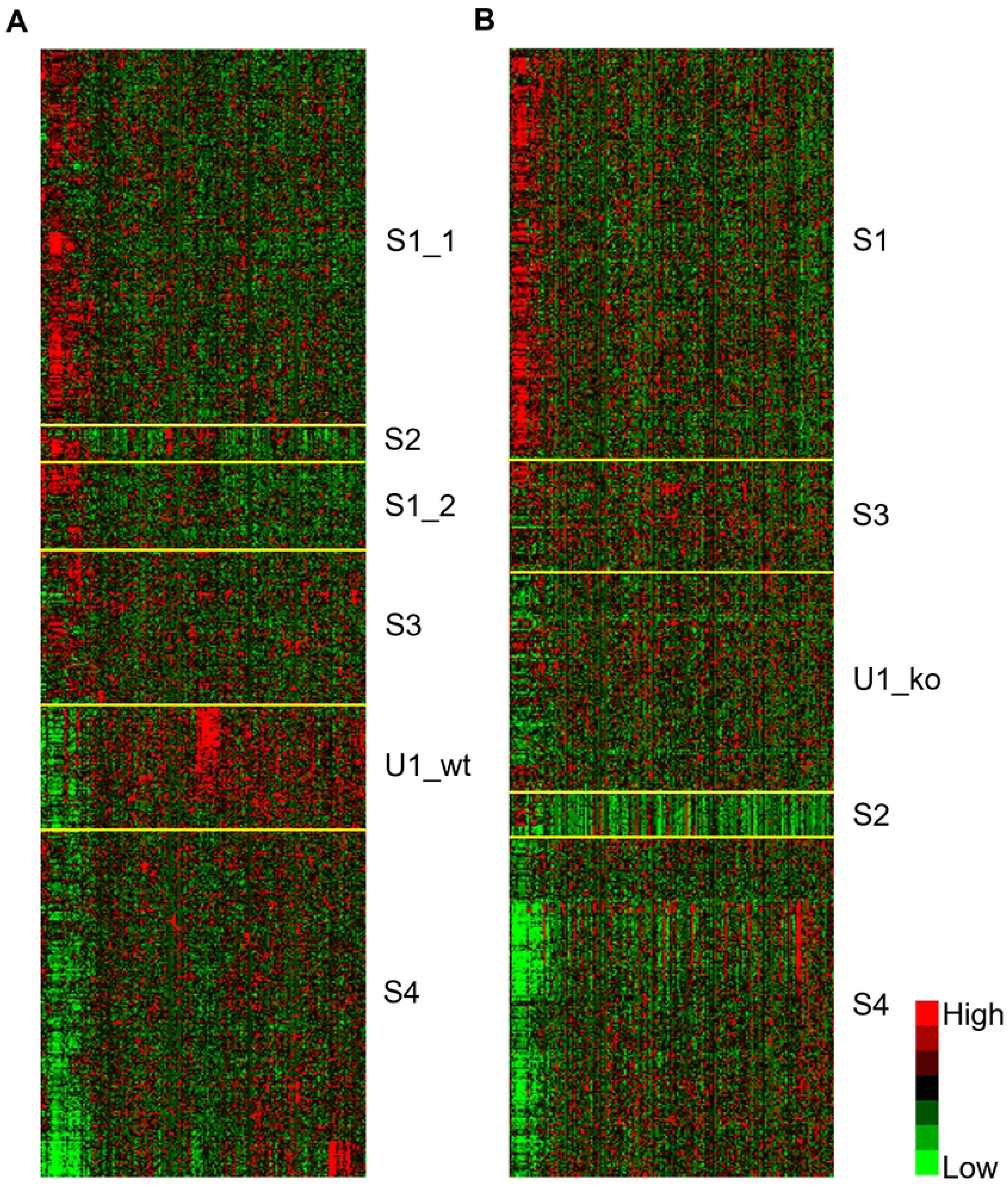
Subclusters of the basal epithelial cells in 5Ht and 6Ho mice revealed by hierarchical clustering based on activities of the common regulons. Heatmap showing the basal epithelial subclusters and regulon patterns of (A) the 5Ht mice and (B) the 6Ho mice. Cells are shown in rows; the 222 common regulons detected in both mouse groups are shown in columns.

We first clustered the basal epithelial cells of the 5Ht mice and their regulons, respectively, using hierarchical clustering (S2A Fig). Then, we clustered the basal epithelial cells of the 6Ho mice, also with hierarchical clustering. Finally, in order to visualize the patterns of the common regulons in the two mouse groups, we reordered the regulons in the 6Ho mice, without any further clustering, in the same order as the clustered regulons in the 5Ht mice (S2B Fig, also see Materials and Methods).

After reordering the basal epithelial cells and their regulons in the 5Ht and 6Ho mice, respectively, using this strategy, several subclusters in the two mouse groups displayed similar regulon activity patterns as clearly shown in Fig 2 and S2 Fig, and could therefore be easily aligned. As such, we identified subclusters S1-S4 and their counterparts in the 5Ht and 6Ho mice. We note that a group of cells within subcluster S1 of the 5Ht mice shared similar regulatory patterns with subcluster S2 of the 6Ho mice; therefore, we designated this group as the counterpart of subcluster S2 of the 5Ht mice. Such clustering behavior is not unusual with hierarchical clustering, as this approach can sometimes generate fragmented clusters with scRNA-seq data (23). Despite hierarchical clustering not being ideal for single-cell partitioning, we found it particularly useful for subcluster alignment, as it can reveal patterns in the clustered data in a hierarchical manner and thus enable us to detect subtle patterns from even a small subcluster of cells. As shown below, we validated these “similar” subclusters in the two mouse groups using UMAP plots. Considering that single-cell data is noisy and that network inference in SCENIC is implemented using a stochastic algorithm, this result was reassuring, indicating that the activities of the regulons detected in the control and the knockout mice are stable and have reproducible patterns.

Other than the “similar” subclusters of the two mouse groups, it was also evident that one subcluster, U1_wt, of the 5Ht mice did not have any counterpart subcluster with a similar regulon pattern in the 6Ho mice, suggesting that knockout of P4HA1 from basal epithelial cells had a direct impact on this subcluster of the control mice. Similarly, one subcluster of the 6Ho mice, U1_ko, had no matching subcluster with a similar regulon pattern in the 5Ht mice, suggesting that this subcluster reflects the knockout effect of P4HA1 in the 6Ho mice.

Next, we examined the distributions of these basal epithelial subclusters in the 5Ht and 6Ho mice using UMAP plots. As shown in Fig 3, each “similar” subcluster and its counterpart in the two mouse groups occupy similar spaces in the UMAP plots. Since these plots were generated based on expression values of all the genes in the mammary cells, and the subclusters of the basal epithelial cells were identified (and also visualized) based on the regulon activities of the same cells, these results indicated that the identified subclusters were valid, not resulting from technical artifacts, and also that each “similar” subcluster of the 5Ht cells and its counterpart in 6Ho were indeed driven by similar regulatory mechanisms.

**Fig 3:**
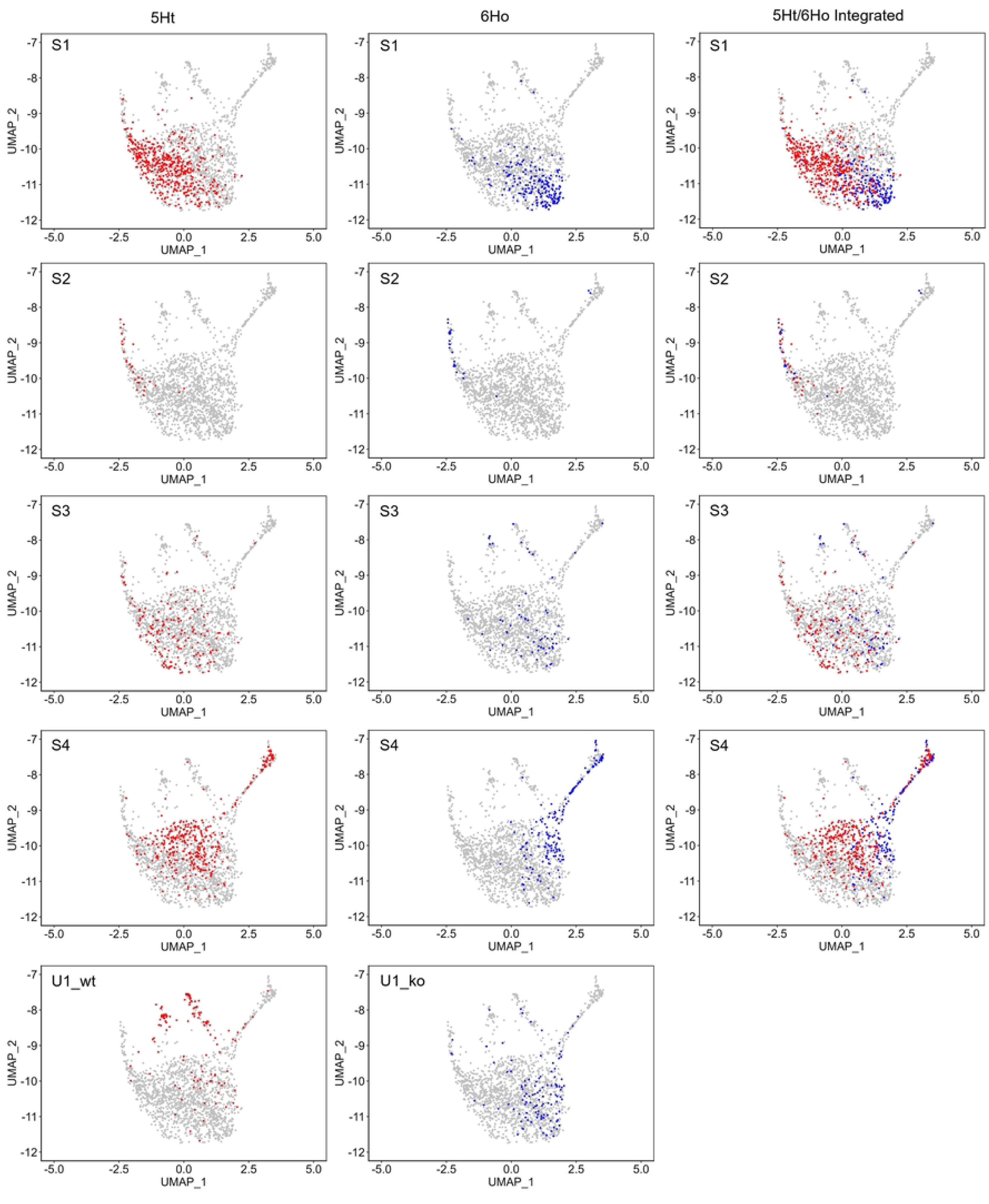
UMAP plots showing all subclusters of the basal epithelial cells in the 5Ht and 6Ho mice. The subclusters were identified based on activities of the common regulons in the 5Ht (left panel) and 6Ho (middle panel) basal epithelial cells. Integrated plots illustrating “similar” subclusters from both mouse groups are in the right panel.

Notably, Fig 3 also shows that subcluster U1_wt of the 5Ht mice and subcluster U1_ko of the 6Ho mice show similar distributions in the UMAP plots, suggesting that they were counterpart subclusters in these mice. However, since our subcluster alignment strategy was based on visualization, we considered U1_wt and U1_ko as the “unique” subclusters (i.e., they do not have any counterpart subclusters in the corresponding mouse group) as they had distinct regulon activity patterns.

### Validation of our results

Since the SCENIC framework has a stochastic component, it was crucial to ensure that the network results we generated for the 5Ht and 6Ho mice were accurate and reliable. To this end, we validated the TF-driven regulons detected in these mice using the Mouse Genome Informatics (MGI) database (24).

We first curated a set of 1032 TFs active in mouse mammary glands using the MGI database. Then, we compared the regulons identified in the basal epithelial cells of the 5Ht and 6Ho mice to this set of TFs. The results are summarized in S2 Table. Of the 268 TF-driven regulons detected in the 5Ht basal epithelial cells, 238 TFs (∼ 89%) overlapped with the MGI set. Similarly, of the 270 TF-driven regulons detected in the 6Ho basal epithelial cells, 239 TFs (∼ 89%) overlapped with the MGI set. We also examined the TFs regulating the unique and common regulons of each mouse group to see how they overlap with the MGI set. A high overlap with the MGI set was observed for these TFs in both mouse groups (S2 Table).

On the other hand, among the 183 TFs included in the network analysis but not detected as regulons in the 5Ht cells, only 83 (∼45%) were part of the MGI set. Similarly, of the 181 TFs not detected as regulons in the 6Ho cells, only 82 (∼45%) were in the MGI set. These findings indicate that the TF-driven regulons identified using SCENIC are biologically valid, aligning with TFs known to be active in mouse mammary gland cells as curated in the MGI database.

### Comparing basal epithelial subclusters in the 5Ht and 6Ho mice

Next, we compared the basal epithelial subclusters of the two mouse groups using differential gene expression (DGE) analysis. For each “similar” subcluster, differentially expressed genes (DEGs) were identified between the 5Ht and 6Ho basal epithelial cells of that subcluster, whereas, for each “unique” subcluster, DEGs were identified between the basal epithelial cells of that subcluster and the remaining basal epithelial cells of the same mouse group. Amongst all the subclusters (“similar” and “unique”), we found that subcluster S4 had the highest number of DEGs, while the “unique” subcluster of the 6Ho mice (U1_ko) had the least. The low number of DEGs in subcluster U1_ko of the 6Ho mice may reflect the effect of the P4HA1-knockout in these mice, i.e., expression levels of the genes in this subcluster become normalized compared to the remaining basal epithelial cells of the same mice.

To further identify differences across the 5Ht and 6Ho mice, we quantified the activity of each regulon within individual cells of these mice using the AUC scores with SCENIC. This score assesses how strongly a TF-driven regulon is active in each cell. As summarized in S3 Table, we found that subcluster U1_wt of the 5Ht mice and subcluster S2 of both mouse groups had the highest numbers of significantly different regulons amongst all the subclusters. On the other hand, subcluster U1_ko of the 6Ho mice had no significantly different regulons.

#### Current findings validated by our previous experimental results

Previously, our experimental results showed that upregulation of P4HA1 in breast cancer cells increased HIF-1 protein stability, subsequently enhancing the stemness of the cells and conferred chemoresistance among breast cancer patients (5). In light of these previous findings, we found two subclusters of the 5Ht mice – U1_wt and S2 – particularly interesting. Our results showed that genes involved in HIF-1 signaling and stem cell differentiation and proliferation were significantly enriched among DEGs in these two basal epithelial subclusters (S4A Table, S5 Table).

In particular, in subcluster S2, our results showed that all 10 genes involved in HIF-1 signaling were downregulated in the 6Ho mice compared to the 5Ht mice (S5 Table, S3 Fig), among which, Hif1a and Vegfa are known to play critical roles in HIF-1 signaling (5, 25). Similarly, 19 DEGs were enriched in stem cell differentiation (S5 Table, S4 Fig), of which 15, including Hif1a and Stat3, were downregulated in the 6Ho mice. These results were consistent with our previous experimental findings, showing that downregulation of P4HA1 led to decreased protein levels of HIF-1, which in turn reduced the stemness of the cells.

Similarly, we found that all 12 DEGs involved in HIF-1 signaling and all 18 genes in stem cell differentiation were upregulated in subcluster U1_wt of the 5Ht mice (S4A Table, Fig 4A, Fig 4B), suggesting that HIF-1 signaling and the stem cell development pathways were suppressed in the basal epithelial cells of the 6Ho mice, explaining why there was no counterpart of subcluster U1_wt with a similar regulon activity pattern in these mice. Also, all 6 DEGs involved in antifolate (which is a chemotherapeutic agent) resistance were upregulated in subcluster U1_wt of the 5Ht mice (S4A Table, Fig 4C). Taken together, these results from subcluster U1_wt of the 5Ht mice and subcluster S2 of both mouse groups agreed with our previous findings showing that downregulation of P4HA1 led to decreased HIF-1 protein levels, reduced stemness of cells, and decreased chemoresistance, thus confirming the validity of our computational findings in this work.

**Fig 4:**
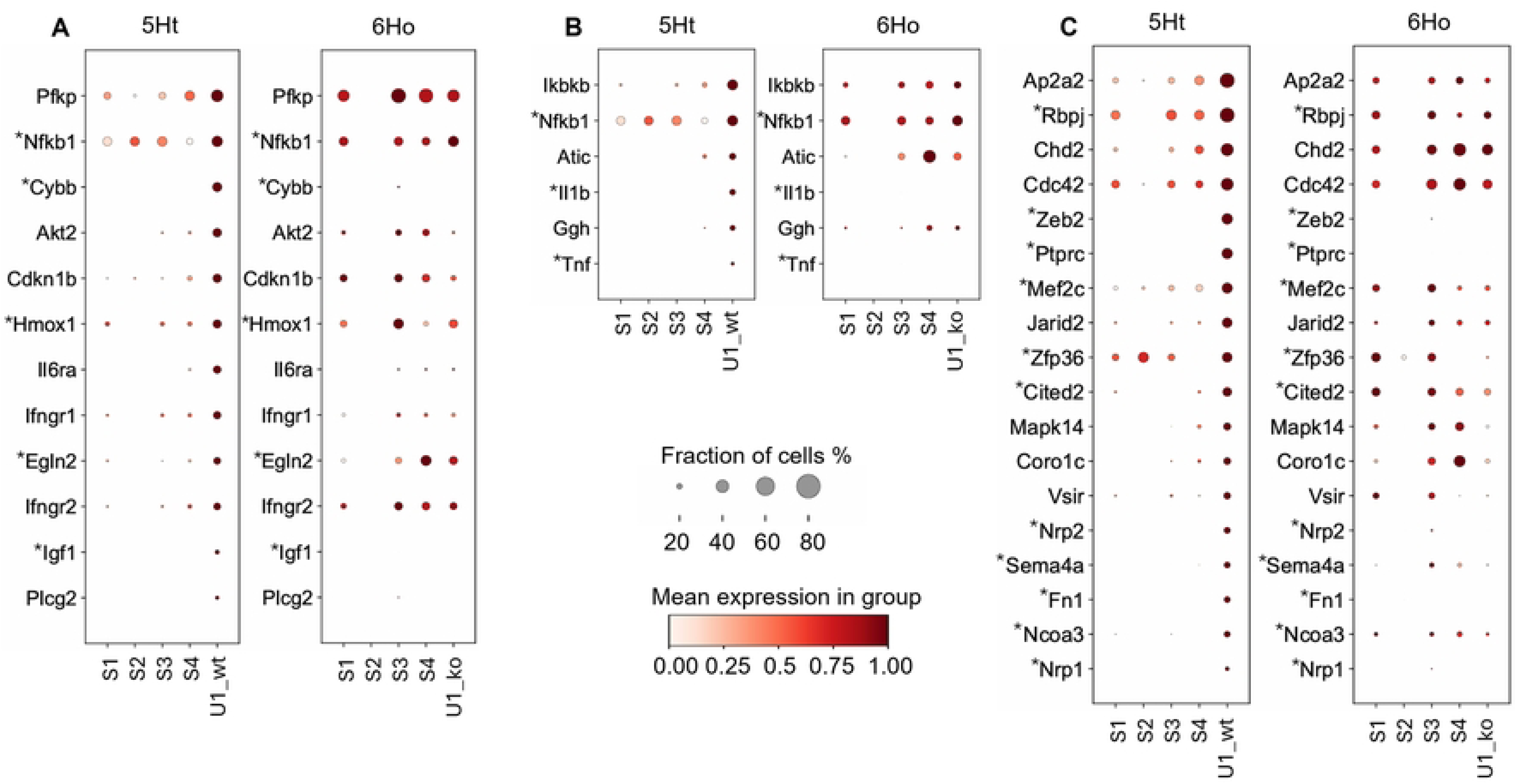
DEGs between subcluster U1_wt and the other 5Ht basal epithelial cells enriched in key signaling pathways. Dot plots display the DEGs enriched in (A) HIF-1 signaling, (B) antifolate resistance, and (C) stem cell development pathways. * indicates DEGs that are either TFs regulating significantly different regulons or their DETGs. DETGs are sorted in descending order based on the fraction of subcluster U1_wt cells in which they are expressed.

#### Novel findings in the current study

Our results also provided novel insights into the molecular mechanisms underlying the mammary basal epithelial cells in the control and the P4HA1-knockout mice. We sought to identify the regulons that played regulatory roles in key signaling pathways in subcluster U1_wt of the 5Ht mice and subcluster S2 of the two mouse groups. In particular, we focused on DETGs, which are target genes of the TF within a regulon that are differential expressed between the corresponding subclusters of the 5Ht and 6Ho mice. We reasoned that it is important to do so because they drive distinct regulatory activities of each regulon in the two mouse groups. This sheds light on how these biological processes are regulated in the cells of these subclusters.

#### Subcluster U1_wt of the 5Ht mice

##### HIF-1 signaling

We did not identify any significantly different regulons that regulated differentially expressed target genes (DETGs) enriched in HIF-1 signaling. This was not unexpected because several key TFs involved in HIF-1 signaling, including Hif1a, Vegfa, and Stat3, were excluded from our network analysis due to pre-processing gene filters set for network reconstruction.

##### Stem cell development

We identified 10 significantly different regulons in this subcluster regulating a shared set of 11 DETGs enriched in stem cell development (S6 Table). These regulons were regulated by the following TFs: Ikzf1, Fli1, Rel, Mafb, Junb, Egr1, Jun, Fos, Fosb, and Elk3. This result agreed with known evidence showing that multiple TFs participating in this pathway and their highly overlapping target genes were synergistically regulated to ensure robust signals for stem cell development (26, 27).

In particular, Fli1, Ikzf1, Rel, and Mafb showed more than a 2-fold upregulation in this subcluster. Dot plots and feature plots in Fig 5 and Fig 6 show the expression of these TFs and their DETGs across all basal epithelial subclusters of the 5Ht and 6Ho mice. It can be seen that their expression levels are upregulated in subcluster U1_wt compared to those in the other subclusters of the two mouse groups.

**Fig 5:**
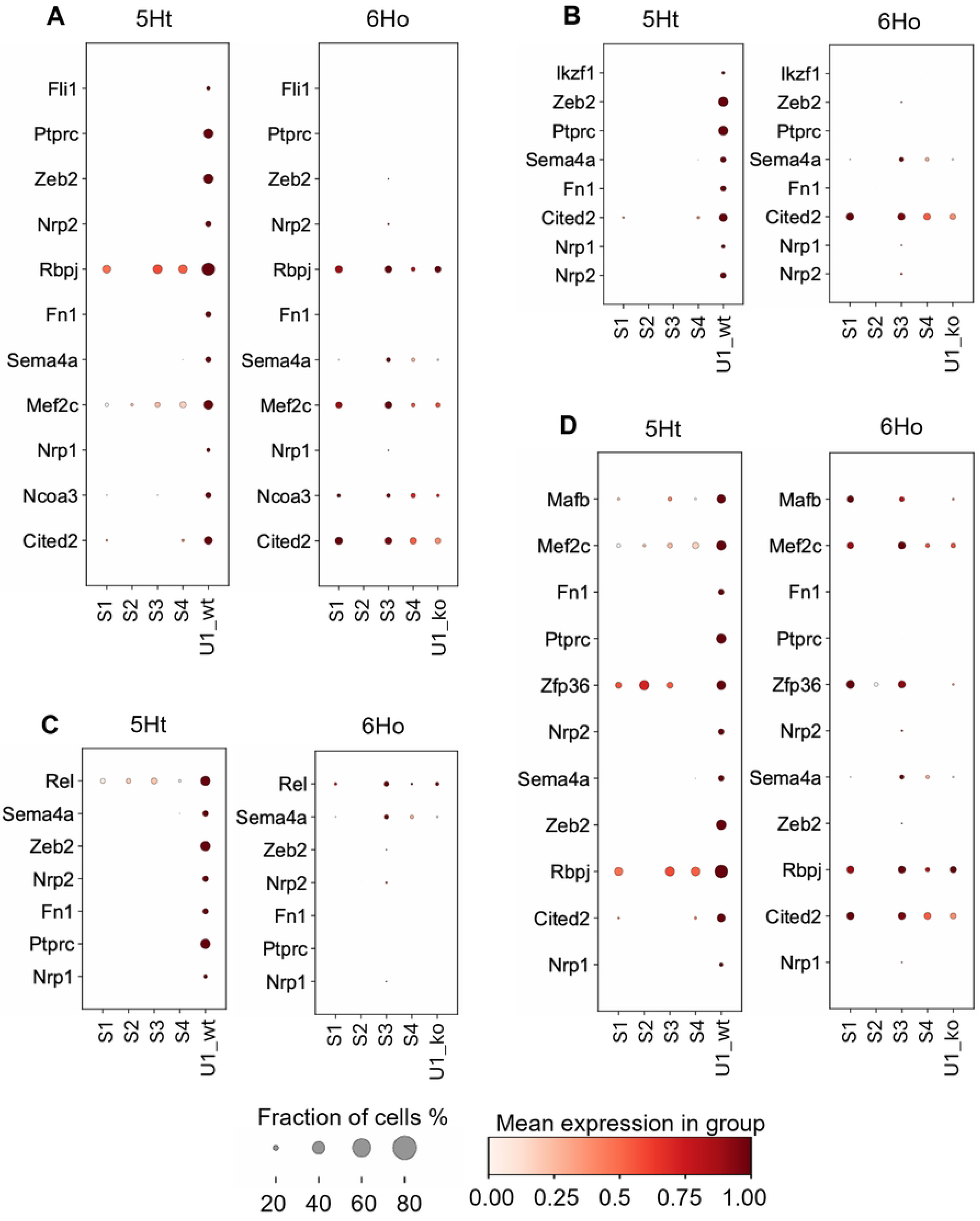
Regulons detected in subcluster U1_wt of the 5Ht mice with the DETGs enriched in the stem cell development pathways. Dot plots show paired comparisons of the (A) Fli1, (B) Ikzf1, (C) Rel, and (D) Mafb regulons and their DETGs across the basal epithelial subclusters of 5Ht and 6Ho mice. Panels (A, B) highlight the DETGs enriched in stem cell development, while (C, D) show the DETGs enriched in stem cell differentiation. DETGs are sorted by their ranks within their respective 5Ht regulon.

**Fig 6:**
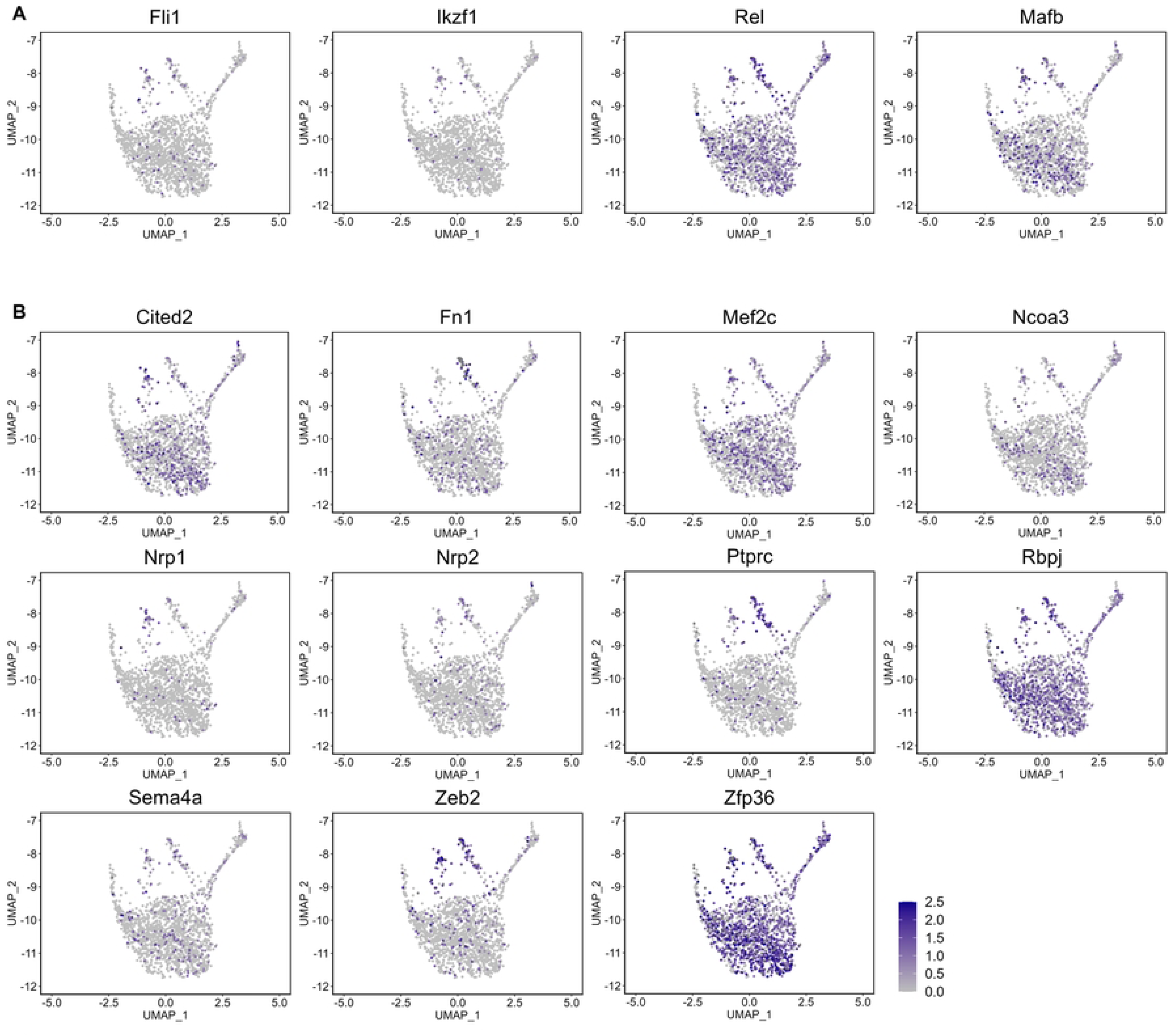
Feature plots of TFs and their overlapping DETGs enriched in stem cell development detected in basal epithelial subcluster U1_wt of the 5Ht mice. Expression levels of (A) TFs regulating significantly different regulons and (B) 11 overlapping DETGs detected in cells of subcluster U1_wt are shown across all basal epithelial cells of the 5Ht and 6Ho mice.

##### Inflammatory and immune responses

Our results show that subcluster U1_wt was significantly enriched with many DEGs involved in numerous inflammatory and immune response processes, e.g., regulation of inflammatory response, acute inflammatory response, leukocyte, B cell, and T cell-mediated immunity, and macrophage activation (S4B Table), all of which, interestingly, have been implicated in crosstalks between inflammation and stem cells (15–17). Moreover, all of these DEGs were upregulated in this subcluster, with the majority having more than a 2-fold change compared to the other basal epithelial cells of the same mice. These results also suggested interactions between basal epithelial cells from subcluster U1_wt of the 5Ht mice and the immune cells such as macrophages and T and B cells.

Using SCENIC, we identified 30 regulons regulating DETGs that were significantly enriched in various inflammatory and immune response processes (S7 Table). The DETGs across these regulons highly overlapped, similar to what we observed with the regulons participating in stem cell development. Notably, 8 of the TFs regulating these 30 regulons – Fli1, Rel, Ikzf1, Mafb, Junb, Egr1, Fos, and Elk3 – were also active in stem cell development (S6 Table, Fig 5, Fig 7), further supporting the notion that there are crosstalks between inflammatory and immune responses and stem cells in this subcluster of the basal epithelial cells and that these 8 TFs play essential roles in such crosstalks.

**Fig 7:**
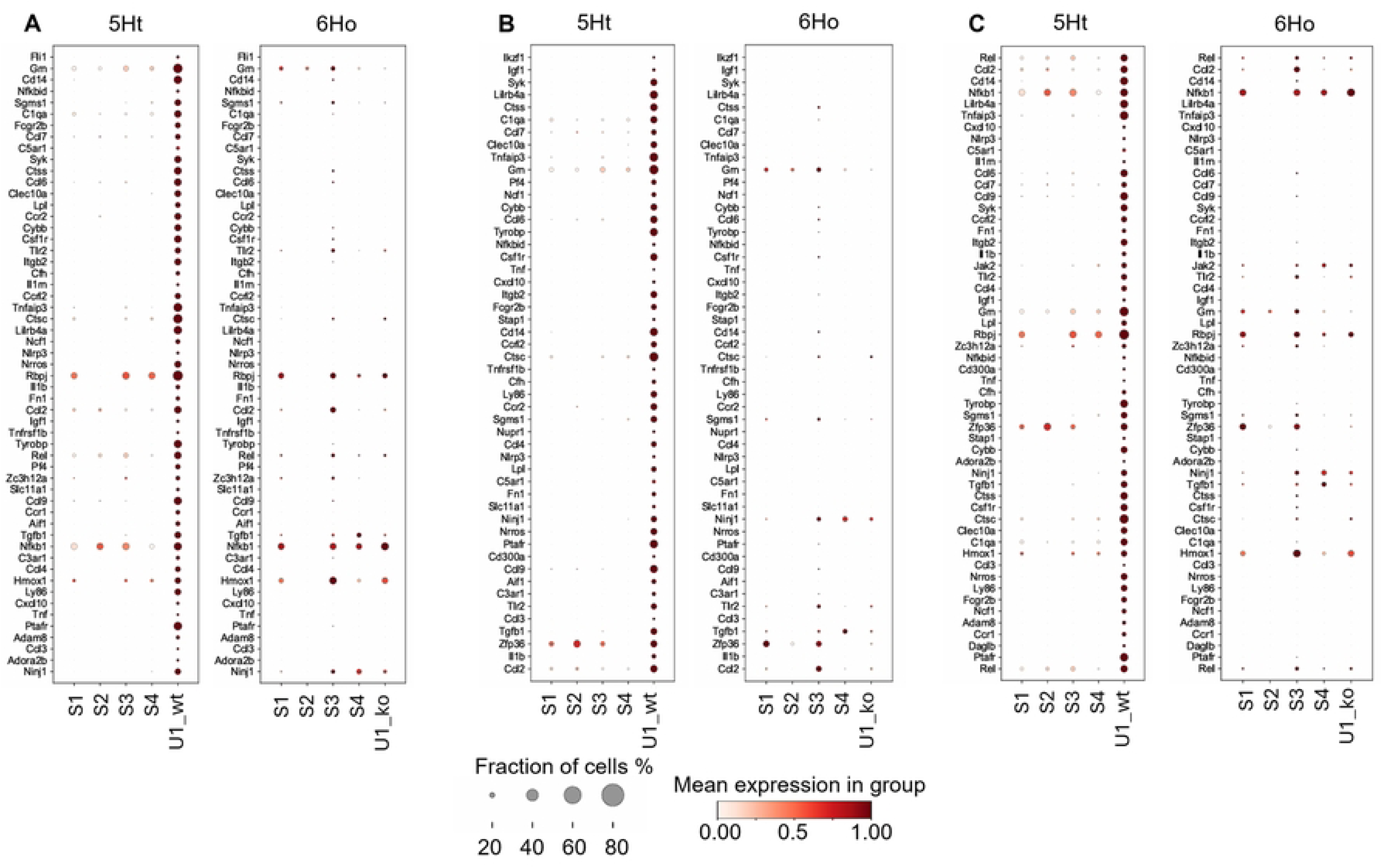
Regulons detected in subcluster U1_wt of the 5Ht mice with the DETGs enriched in the inflammatory response pathway. Dot plots show paired comparisons of the (A) Fli1, (B) Ikzf1, and (C) Rel regulons and their DETGs across the basal epithelial subclusters of 5Ht and 6Ho mice. Panels (A-C) show the DETGs enriched in inflammatory response. DETGs are sorted by their ranks within their respective 5Ht regulon.

In particular, of all the 30 regulons, Fli1 regulated 54 DETGs involved in the inflammatory response pathway, all of which were upregulated in subcluster U1_wt. Moreover, 51 DETGs had more than a 2-fold change compared to the other 5Ht basal epithelial cells (Fig 7A). Feature plots in Fig 8 show the distributions of 8 of these DETGs across all basal epithelial cells of the 5Ht and 6Ho mice; it can be seen that their expression levels align very well with the expression profile of Fli1 in the 5Ht subcluster U1_wt cells (also see Fig 3 for the distribution of the 5Ht subcluster U1_wt cells in the UMAP plot). Following our findings, a comprehensive literature review highlighted Fli1 as a key proto-oncogene with crucial roles in hematopoiesis and vascular development (28). Fli1 is also recognized as one of the top regulators of blood stem and progenitor cells (29). Its overexpression has been implicated in highly malignant triple-negative breast cancer, where it promotes tumor progression by driving proliferation, inhibiting differentiation, and enhancing cellular survival under hypoxic conditions (30–32). All of this evidence supports our findings.

**Fig 8:**
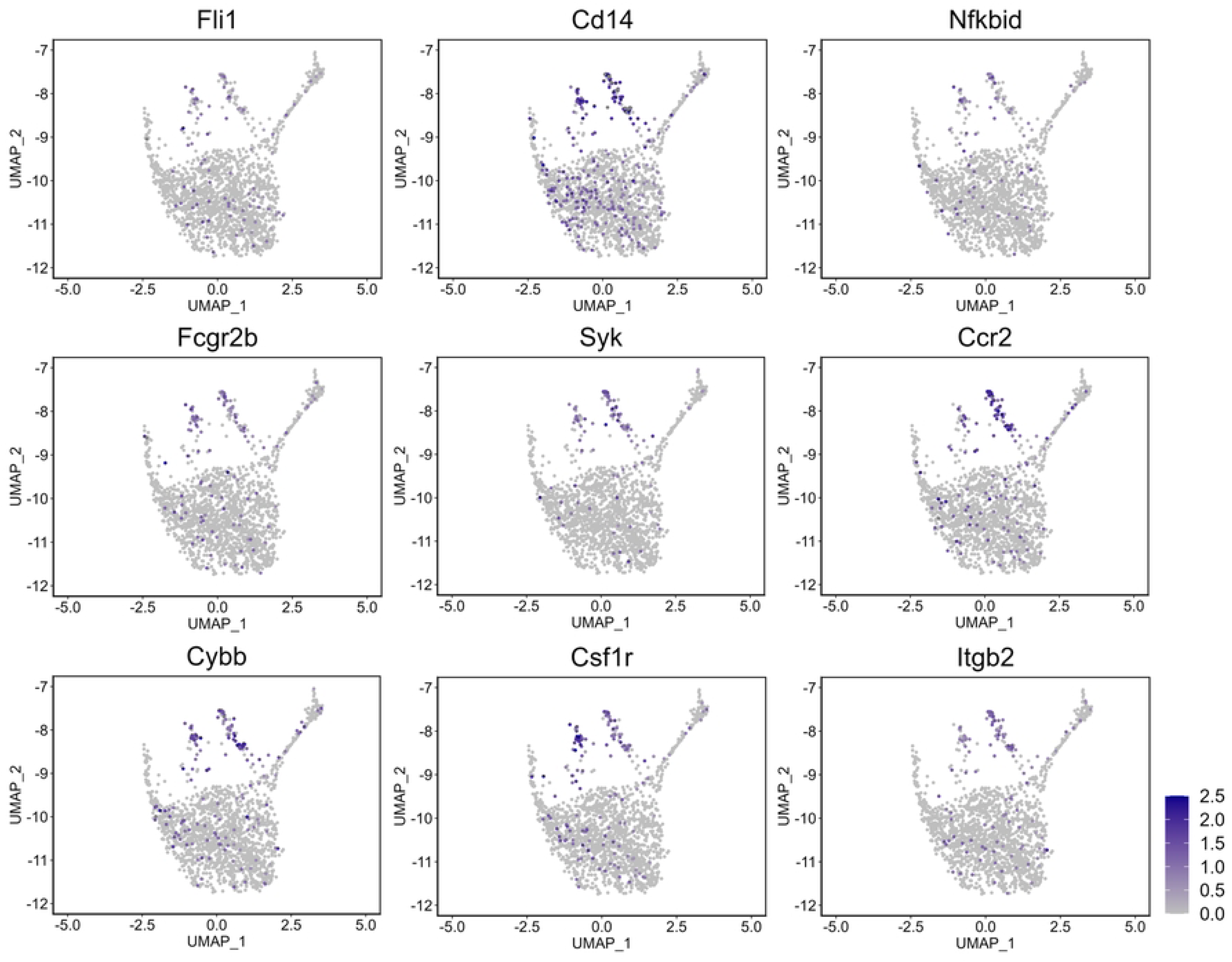
Feature plots of the Fli1 regulon and its DETGs enriched in the inflammation response pathway detected in basal epithelial subcluster U1_wt of the 5Ht mice. Expression levels of Fli1 (TF) and several DETGs (within the regulon) detected in cells of subcluster U1_wt are shown across all basal epithelial cells of the 5Ht and 6Ho mice.

Taken together, these results indicated that there are crosstalks between the inflammatory and immune responses and stem cell development pathways in subcluster U1_wt of the 5Ht mice, which are mediated by multiple overlapping TFs regulating both processes. Moreover, these TFs regulated a large number of overlapping DETGs to provide robust regulatory signals for each of the biological processes. Furthermore, these results suggested that knockout of P4HA1 leads to reduced stemness and inflammation in this subcluster of the cells in the control mice.

#### Subcluster S2 of the 5Ht and 6Ho mice

Subcluster S2 had similar regulon activity patterns in the 5Ht and 6Ho mice, suggesting that knockout of P4HA1 did not significantly affect these cells.

##### HIF-1 signaling

Even though we detected DEGs such as Hif1a, Vegfa, and Stat3 enriched in HIF-signaling in this subcluster of the cells, we did not identify any significantly different regulons participating in HIF-1 signaling between the 5Ht and 6Ho mice due to the same reason stated earlier for subcluster U1_wt of the 5Ht mice, where these key TFs were excluded from the analysis due to gene filtering criteria.

##### Stem cell development

We identified 2 regulons in subcluster S2 of the 5Ht mice and 7 in subcluster S2 of the 6Ho mice with the DETGs significantly enriched in stem cell differentiation and development (S8 Table, S5 Fig, S6 Fig). We noticed that the Egr1 and Junb regulons were detected in subcluster S2 of both the mouse groups, with nearly identical DETGs enriched in stem cell development (S5A Fig and S6A Fig; S5B Fig and S6B Fig); these consistent results indicated that downregulation of these TFs in the 6Ho mice led to downregulation of the DETGs participating in stem cell development in these cells (S6A Fig, S6B Fig).

For the 6Ho mice, we identified 5 additional regulons that have DETGs enriched in stem cell differentiation, including Rel, Irf1, Elf1, Maf, and Klf4 (S8B Table). In particular, 4 TFs, Rel, Irf1, Elf1, and Maf, showed more than a 2-fold downregulation in subcluster S2 of the 6Ho mice, which subsequently led to the downregulation of the corresponding DETGs that are enriched in the stem cell development pathway (S6 Fig).

Unlike subcluster U1_wt of the 5Ht mice, we did not detect any DETGs in subcluster S2 of the two mouse groups that were enriched in the inflammatory and immune response pathways. Overall, these results indicated that knockout of P4HA1 had significant effects on reducing the stemness of the cells in the basal epithelial subcluster S2 of the 6Ho mice mediated by transcriptional regulons.

### Experimental validation

Our results suggest crosstalks between stem cell development and inflammatory signaling in basal epithelial subcluster U1_wt of the 5Ht mice. Specifically, we found that 21 DEGs participating in macrophage activation were upregulated in subcluster U1_wt of the 5Ht mice (S4 Table). Notably, several of these 21 genes, e.g., Tyrobp, Syk, Csf1r, and Tnf, were the DETGs regulated by multiple TFs, including Fli1, Rel, Ikzf1, and Mafb (S7 Table), which we also identified as the key regulators orchestrating both stem cell development and inflammatory signaling in the subcluster U1_wt cells. Given the crucial function of macrophages in inflammatory response as well as recent evidence suggesting macrophages promote breast cancer initiation in premalignant mammary lesions (33), we performed immunohistochemistry (IHC) to examine macrophage accumulation in the mammary glands of the 5Ht and 6Ho mice.

To identify macrophages, we immunostained sections of the mouse mammary gland with F4/80 antibodies (34). These antibodies recognize and bind to the F4/80 antigen, a 160-kDa glycoprotein belonging to the EGF-TM7 family, expressed at high levels on the surface of macrophages (see Materials and Methods for additional details).

As shown in Fig 9, macrophage presence was notably reduced in the 6Ho mammary tissue compared to that of the 5Ht mouse. This suggests that the macrophage-driven inflammatory response is suppressed in the mammary glands of the P4HA1-deficient 6Ho mice. These findings validate our results, confirming that macrophage activation is indeed disrupted in the P4HA1-deficient model, thus reinforcing the critical role of P4HA1 in supporting macrophage-mediated inflammatory processes in mammary tissue.

**Fig 9:**
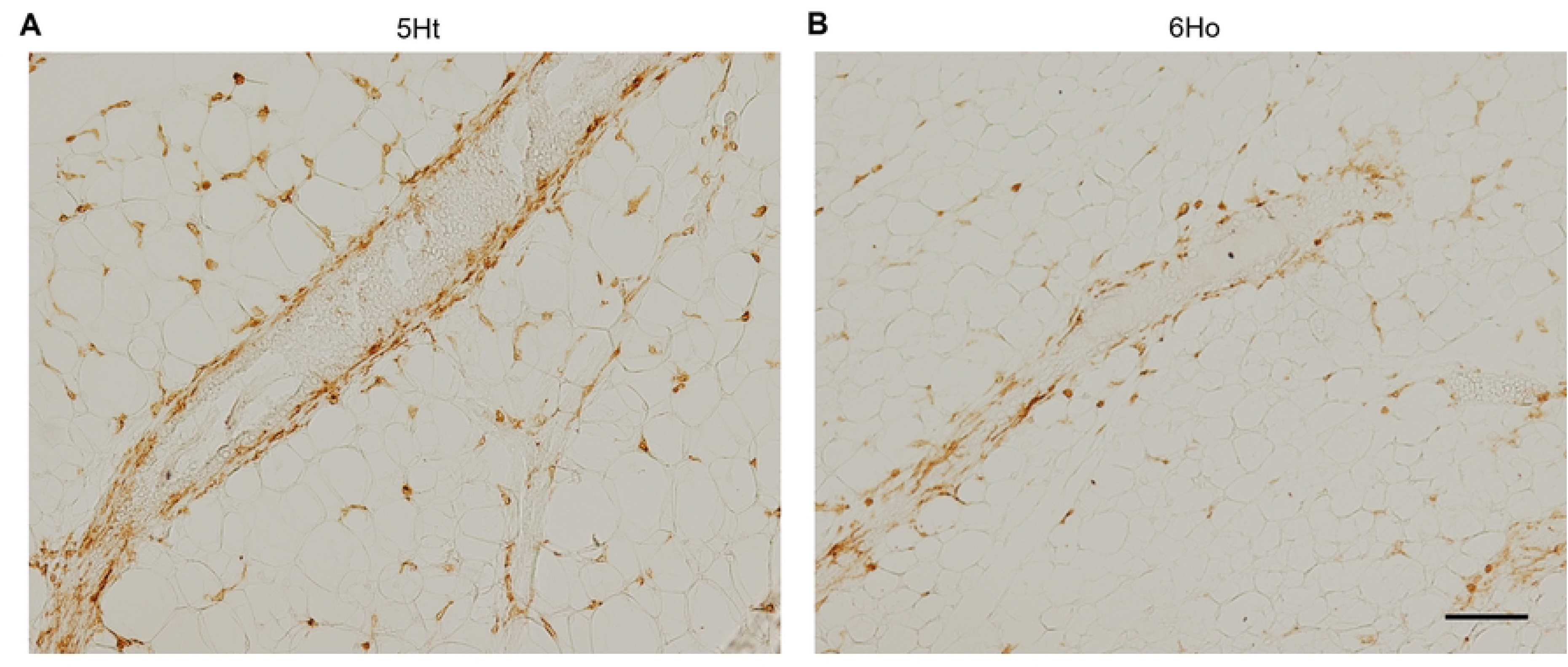
Immunostaining of macrophages in the mammary glands of the 5Ht and 6Ho mice. Macrophages were identified by F4/80 antibody, shown by brown staining. (A) The mammary tissue of the 5Ht mouse shows a higher presence of macrophages, which may indicate more active immune involvement. (B) In contrast, the mammary tissue of the 6Ho mouse has fewer macrophages, suggesting that the P4HA1 deficiency could be linked to a reduction in macrophage-driven inflammation in these mice. Scale bar, 100 µm.

## Discussion

In this study, we investigated the molecular mechanisms of P4HA1 in mammary basal epithelial cells of control (5Ht) and P4HA1-knockout (6Ho) mice using scRNA-seq data. Gene network analysis revealed distinct regulatory differences between the basal epithelial cells of the two mouse groups, with the absence of P4HA1 altering important pathways, including HIF-1α signaling and stem cell development, in some subclusters of these cells. Leveraging the SCENIC pipeline, we identified TF-driven regulons activated in mammary basal epithelial cells of both mouse groups. Furthermore, we focused on analyzing DETGs of the detected regulons in the two mouse groups since they drive distinct molecular activities of regulons in these mice. Similar to focusing on DEGs when comparing gene expression profiles, analyzing DETGs within regulons provides targeted insights into the cellular mechanisms that differentiate different regulatory networks. By doing so, our results demonstrated the presence of complex crosstalks between stem cell development and inflammation in a basal epithelial subcluster of the control mice, providing novel insights into how the absence of P4HA1 affects these biological processes.

One of the significant challenges in single-cell network analysis is the inherent noise and technical variability associated with the scRNA-seq data. This problem is compounded when comparing networks from different biological conditions, such as the 5Ht and 6Ho mouse groups, where distinguishing genuine biological differences from technical artifacts becomes critical. To address these challenges, we employed several validation strategies throughout the analysis. First, we cross-validated our results in the control mice with previously published studies, ensuring that our findings aligned with known biological knowledge. Second, we carefully aligned subclusters of the basal epithelial cells in both 5Ht and 6Ho mice using reproducible regulon activity patterns (Fig 2) and confirmed their similarity in the UMAP plots (Fig 3). This approach enabled us to identify subclusters governed by the shared regulatory mechanisms, ensuring that the observed patterns reflected genuine biological processes rather than artifacts introduced by noise or stochastic variability. Third, we validated the TFs of the regulons against curated public databases, such as the MGI database, focusing on TFs known to be active in the mouse mammary gland. We did so to ensure that the detected TFs and their regulons were biologically valid.

Furthermore, we performed experimental validation using immunohistochemistry to assess the macrophage accumulation in the mammary glands of the two mouse groups. We observed a significant reduction in macrophage presence in the 6Ho mammary tissue compared to the 5Ht mouse (Fig 9). This further confirmed that the absence of P4HA1 suppresses macrophage activation in the knockout mice. Together, these validation strategies ensured that the reconstructed networks and our findings were robust, reliable, and reflective of genuine biological differences between the 5Ht and 6Ho groups, mitigating concerns about noise and technical artifacts resulting from the scRNA-seq data.

Our results revealed several novel findings, particularly in identifying crosstalks between stem cell development and inflammatory and immune responses in subcluster U1_wt of the basal epithelial cells of the control mice. We found that many TFs regulated overlapping gene sets involved in both processes. Notably, 8 TFs—including Fli1, Rel, Ikzf1, and Mafb—emerged as key regulators orchestrating stem cell maintenance and inflammatory signaling. Building on our findings, a literature review further emphasized the critical functions of Fli1 as a proto-oncogene. It is notably one of the ten key regulators of blood stem and progenitor cells, essential for maintaining vascular integrity and promoting blood vessel formation (28, 29). Additionally, the upregulation of Fli1, observed in aggressive triple-negative breast cancer (TNBC) cell lines (32) reinforces its potential role in driving cancer progression. Similarly, Rel, a key member of the NF-κB family, is predominantly involved in immune regulation, and it is frequently amplified or mutated in lymphoid malignancies (35). Ikzf1 is essential for lymphocyte development and homeostasis (36). Finally, Mafb is involved in macrophage differentiation and immune response modulation. Dysregulation of MAF family TFs, including Mafb, has been associated with cancer proliferation, metastasis, and the regulation of tumor-associated macrophages (TAMs) (37).

Although crosstalks between stem cell development and immune cells were found in the control mice in our study, there is substantial evidence from various cancer studies that stem cells, and immune cells interact closely in the tumor microenvironment, influencing tumor progression and immune evasion. For example, NF-κB activation (also shown in our results, S7 Table) contributes to immune regulation and plays a significant role in the proliferation and differentiation of stem cells in various cancers, including breast cancer (38, 39). Thus, these crosstalks between stem cells and immune cells have been found to support tumor growth, metastasis, and drug resistance (16).

Our study also highlighted the redundant functions of several TFs in regulating stem cell development and inflammatory responses. We observed that multiple TFs (such as Fli1, Ikzf1, and Mafb) regulated shared target genes, creating a highly redundant regulatory network. Evidence has shown that functional redundancy is often critical for maintaining cellular stability, particularly in dynamic environments such as stem cell development (26, 40) and tumorigenesis (41); it allows for compensatory regulation when individual TFs are disrupted, ensuring that critical processes, such as immune responses and stem cell differentiation, continue unhindered (26).

By leveraging single-cell network inference techniques and validating our results through multiple strategies, we provide novel insights into the molecular mechanisms of P4HA1 in potentially regulating both stem cell differentiation and inflammation in the mouse mammary gland. The absence of the key subcluster of basal epithelial cells in P4HA1-deficient 6Ho mice highlights how P4HA1 influences the interplay between stem cell development and inflammation. These results suggest that the inhibition of P4HA1 could disrupt the crosstalks between these signaling pathways, highlighting the potential of targeting P4HA1 as a therapeutic strategy for breast cancer treatment.

## Materials and Methods

### Ethics statement

The mouse experiments (Protocol Number: 2020-3639) were approved by the Institutional Animal Care and Use Committee (IACUC) at the University of Kentucky. All mouse experiments were performed with the Division of Laboratory Animal Resources guidelines at the University of Kentucky.

### Mammary gland isolation and dissociation for scRNA-seq data generation

In this study, we generated scRNA-seq data from two groups of mice: K14-cre; P4HA1 ^loxp/+^ (control, 5Ht) mice and K14-cre; P4HA1 ^loxp/loxp^ (P4HA1-knockout, 6Ho) mice. P4HA1 expression was specifically silenced in the basal mammary epithelial cells in 6Ho mice.

To isolate mammary gland cells for scRNA-seq, we dissected mammary glands from both groups (two mice in each group) and mechanically dissociated them. The finely minced tissue was transferred to a digestion mix consisting of DMEM/F12 (Gibco) + 10 mM HEPES (Gibco) + collagenase (Roche) + 200 U ml⁻¹ hyaluronidase (Sigma) + gentamicin (Gibco) for 3 hours at 37°C and vortexed every 30 minutes. After the lysis of red blood cells in NH_4_Cl, cells were briefly digested with warm 0.05% Trypsin-EDTA (Gibco), 5 mg ml^−1^ dispase (Sigma), and 1 mg ml^−1^ DNase (Sigma), and filtered through a cell strainer (BD Biosciences) to remove debris, yielding a single-cell preparation for sequencing.

Propidium iodide (PI, Sigma) was used at a working concentration of 1 μg/ml, prepared by diluting the stock solution 1:1000 to detect dead cells. After sorting, cells were spun down and resuspended. Samples were manually counted using an improved Neubauer chamber, and the cell concentration was normalized. Equal numbers of cells per sample were processed for scRNA library preparation.

### ScRNA-seq data generation and pre-processing

ScRNA-seq libraries for 5Ht and 6Ho mice were generated using the 10X Genomics Chromium platform. The resulting gene expression libraries were mapped to the mouse reference genome (mm10, 10x Genomics pre-built reference mm10-2020-A) using Cell Ranger (v.6.1.1). For each scRNA-seq library, we applied quality control (QC) filters on the Cell Ranger filtered matrix to remove low-quality cells, defined as those with ≤ 200 detected features or ≥ 10% of reads mapping to mitochondrial genes. Additionally, cells with unique molecular identifier (UMI) counts ≥ 30,000 were identified as potential doublets and were excluded from further analysis.

Post-QC, the filtered datasets were integrated using the recommended sequential procedures and functionalities described in the Seurat (v.4.4.0) package. The raw gene expression matrices were normalized using the ***SCTransform*** function, which applies regularized negative binomial regression to mitigate the influence of technical noise. Next, ***SelectIntegrationFeatures*** was used to select integration features, which were then passed to ***PrepSCTIntegration*** to prepare the objects for data integration. Integration anchors were identified with ***FindIntegrationAnchors***, and the datasets were merged using the ***IntegrateData*** function. The integrated data was then clustered and visualized using the UMAP algorithm based on the top 30 principal components (PCs) computed by PCA. For clustering, we identified the 20 nearest neighbors using the ***FindNeighbors*** function and performed clustering using ***FindClusters*** with a resolution of 0.5, resulting in 25 distinct clusters.

Cluster markers were identified using the Wilcoxon rank-sum test by comparing each cluster with the rest of the cells. We performed the cell type identification using known canonical cell markers, such as Epcam, Krt14, and Krt5 for basal epithelial cells, Epcam, Aldh1a3, Gata3, and Cdh1 for epithelial luminal cells, Cd14 and Adgre1 for macrophages, and Cd3d for T cells.

After quality control and data integration, the ***filtered scRNA-seq dataset*** comprised 12,472 cells and 19,140 genes. This dataset was then used for subsequent downstream analyses, including network and differential gene expression analyses, as described below.

### Network Analysis

#### ScRNA-seq data pre-processing

We pre-processed the ***filtered scRNA-seq dataset*** for network analysis using Scanpy (v.1.9.4) (42) and its pre-processing modules. First, we filtered the cells and the genes separately for 5Ht and 6Ho mice using the ***filter_genes*** and ***filter_cells*** functions; we retained the cells expressing fewer than 5000 genes and the genes expressed in fewer than 3 cells. Post-filtering, we had 7,624 cells and 18,433 genes in ***the filtered 5Ht dataset*** and 4,757 cells and 18,052 genes in ***the filtered 6Ho dataset***. Subsequently, we removed the genes with the “Rik” suffix (e.g., 5730419F03Rik) and those starting with “Gm” (e.g., Gm31812) to focus our analysis on well-annotated genes. Also, we combined the ***filtered 5Ht and 6Ho datasets*** to create ***a combined dataset*** that contains data for both groups of mice.

Then, we performed the subsequent processing steps on the filtered 5Ht, 6Ho, and combined datasets separately. We normalized total counts to 10,000 per cell using ***normalize_total*** and natural log-transformed the data. We then identified highly variable genes (HVGs) using the ***highly_variable_genes*** function and retained the HVGs with minimum mean expression of 0.0125, maximum mean expression of 3, and minimum dispersion of 0.45 for subsequent analyses. Finally, the counts were scaled for each gene to have a unit variance and zero mean.

To facilitate comparative analysis, we took the union of the HVGs retained in the 5Ht, 6Ho, and the combined datasets, which resulted in 4,838 genes. Consequently, ***the pre-processed 5Ht dataset*** contained 7624 cells and 4838 genes, while ***the pre-processed 6Ho dataset*** contained 4757 cells and 4838 genes. These datasets were then used for network analysis.

#### SCENIC network analysis

Network analysis was conducted using the SCENIC (v.0.12.1) pipeline in Python, which was applied to basal epithelial cells of the 5Ht and 6Ho mice, respectively. The SCENIC workflow involves three main steps: network inference, motif enrichment, and cellular enrichment of regulons.

##### Network inference

SCENIC utilizes the stochastic gradient boosting machine algorithm ***GRNBoost2*** (43) to infer gene regulatory networks based on the co-expression of TFs and their target genes. A set of 1860 TFs obtained from the cisTarget resources database (https://resources.aertslab.org/cistarget) and ***the pre-processed 5Ht and 6Ho datasets*** were used, respectively, as input for GRNBoost2. Of these 1860 TFs, 451 TFs were present in the pre-processed datasets and were the basis of our network analysis. The output of this algorithm was an edge list mapping TFs to their target genes with an associated importance score obtained from the regression model.

To account for the stochastic variability of GRNBoost2, we ran the algorithm 20 times for each mouse group, ensuring that the associations between the identified TFs and their target genes were stable and reproducible. For results from each run of the algorithm, we performed motif enrichment as described below.

##### Motif enrichment

The network inferred from above was then refined by assessing motif enrichment in the regulatory regions of the identified target genes. This step involved generating modules and regulons. Modules are sets of the identified TFs and their target genes, generated using the ***modules_from_adjacencies*** function. This function filters the TFs and their target genes by selecting the top-ranked target genes for each identified TF based on their importance scores obtained from the tree-based regression model. These modules were then used to generate regulons through motif enrichment using ***prune2df*** and ***df2regulons***.

##### Ranking target genes for each identified regulon

We summarized the results from all 20 runs by calculating summary statistics of the detected target genes for each regulon, including target gene frequencies, importance scores, and the mean, median, and standard deviation of expression levels of the genes. The expression values were calculated across the cells of the particular mouse group. Target genes were ranked based on the frequency of each gene’s occurrence in the regulon across 20 runs of the SCENIC analysis, with a higher frequency indicating a higher rank. They were then sorted by rank within each regulon.

##### Cellular enrichment of regulons

Regulons detected across all 20 runs were aggregated by the union operation for each mouse group to estimate the cellular enrichment of the regulons by the AUC scores using the ***AUCell*** function. The output of this step was an AUC matrix, with cells in rows and the detected regulons in columns. A higher AUC score indicates that more target genes of a regulon are actively expressed, reflecting the activity of the TF in regulating gene expression within that specific cell.

#### Validation of the identified regulons

The MGI database is a collection of expertly curated information and resources that integrates the Mouse Genome Database (MGD) and the Gene Expression Database (GXD) to support experimental and computational investigations of the laboratory mouse genome. We used the MGI database (v.6.23) to generate a set of TFs active in mouse mammary gland cells. To obtain this set, we first identified all genomic features active in mammary gland cells of mice, which resulted in a set of 23,754 features. We then filtered this set only to include protein-coding genes with DNA-binding transcription factor activity. This resulted in a set of 1032 transcription factors we used for validation.

### Cluster Analysis and subcluster alignment

To identify subclusters of basal epithelial cells in 5Ht and 6Ho mice, we first standardized the AUC matrices so that each column (representing a regulon) has a mean of 0 and a standard deviation of 1. Then, we applied agglomerative, complete-linkage hierarchical clustering on the standardized AUC matrices to partition the genes and the cells into distinctive groups based on the cellular activities of regulons for both 5Ht and 6Ho mice. Correlation was used to measure the similarity between variables (regulons or cells). The hierarchical clustering algorithm was implemented using Cluster 3.0 (v.1.59).

Clustering was first performed on the basal epithelial cells and the regulons in the 5Ht mice. To facilitate the alignment of the subclusters in the 5Ht and 6Ho mice, we reordered the regulons in the 6Ho mice to match the order of the clustered 5Ht regulons. We then performed clustering on the basal epithelial cells of the 6Ho mice without clustering the regulons. This strategy facilitated comparison between the two mouse groups by maintaining the same regulon order.

The subclusters in the two mouse groups were then aligned based on discernible patterns, as revealed in the heatmaps of the clustered AUC matrices. These subclusters were then categorized as either “similar” (with similar patterns in both mouse groups) or “unique” (with a pattern specific to only one mouse group).

### Differential Gene Expression (DGE) Analysis

For DGE analysis, we used the ***filtered scRNA-seq dataset*** containing 12,472 cells and 19,140 genes from both 5Ht and 6Ht mice (see the “ScRNA-seq data generation and pre-processing” subsection in “Materials and Methods” for details). Then, we excluded cells expressing more than 5,000 genes and discarded genes expressed in fewer than 3 cells. This resulted in a dataset containing 12,381 cells and 19,032 genes. We then normalized and natural log-transformed the data using *Seurat’s* (v.5.1.0) ***NormalizeData*** function. DEGs were identified using the ***FindMarkers*** function with the MAST test and an absolute fold change threshold of 1.5. We also set the minimum difference in the fraction of detection (“min.diff.pct”) between the two compared groups and the “min.pct” to be 0.15, respectively. We used the MAST test because it identifies DEGs between two groups of cells using a hurdle model tailored to scRNA-seq data (44). ***FindMarkers*** used the Bonferroni correction to account for multiple hypothesis testing.

For the “similar” subclusters (S1-S4), DEGs were identified between each subcluster and its counterpart in the two mouse groups. For each “unique” subcluster (i.e., U1_wt of the 5Ht mice or U1_ko of the 6Ho mice), DEGs were identified between the “unique” subcluster and all the other basal epithelial cells of the same mouse group.

### Identification of significantly different regulons between the 5Ht and 6Ho mice

For each subcluster of basal epithelial cells, we identified regulons with significantly different AUC scores between the two mouse groups using the Wilcoxon rank-sum test, followed by the multiple testing correction using the Benjamini-Hochberg procedure to control the false discovery rate (FDR) (45). For the “similar” subclusters (S1-S4), the AUC scores of each regulon were compared between each subcluster and its counterpart in the two mouse groups. For the “unique” subclusters (i.e., U1_wt of the 5Ht mice and U1_ko of the 6Ho mice), the AUC scores were compared between cells in each “unique” subcluster and all the other basal epithelial cells of the same mouse group. Regulons with an adjusted p-value < 0.05 and an absolute mean AUC score difference > 0.25 were considered significantly different.

### Functional enrichment analysis

To identify functional groups significantly enriched among the DETGs in the detected regulons from the network analysis and the DEGs from the DGE analysis, we performed functional enrichment analysis using the ***clusterProfiler*** package in R (46). This package estimates overrepresentation in Gene Ontology (GO) and KEGG pathways among genes of interest using the hypergeometric test. The set of the 4838 unique genes used for the network analysis was employed as the gene universe for the enrichment analysis of the DETGs. The set of 19,032 unique genes used in the DGE analysis was employed as the gene universe for DEGs. A GO term or a KEGG pathway was considered significantly enriched if it had an adjusted p-value < 0.05, with the multiple testing correction using the Benjamini-Hochberg procedure.

### Visualization

The UMAP plots were generated using the ***DimPlot*** function in Seurat to illustrate the spatial distributions of the identified subclusters in a 2D space. The ***FeaturePlot*** function was used to visualize expression levels of genes across all the basal epithelial cells in the 5Ht or 6Ho mice. Dot plots were generated using the ***dotplot*** function from Scanpy. For heatmaps, we utilized TreeView (v.1.2.0) (https://sourceforge.net/projects/jtreeview/files/jtreeview/1.2.0/) to display the hierarchical clustering of the AUC matrices in the two mouse groups.

### Immunohistochemistry analysis of the mammary glands in 5Ht and 6Ho mice

To identify macrophages, mammary gland sections were immunostained using F4/80 antibodies, a marker known to label macrophages. The sections were deparaffinized and rehydrated through a graded alcohol series (100%, 95%, and 70%) into phosphate-buffered saline (PBS) solution to remove paraffin and restore the tissue to an aqueous state suitable for staining. Endogenous peroxidase activity was blocked by incubation with 3% H₂O₂ for 20 minutes to prevent non-specific background staining. For antigen retrieval, slides were steamed in citrate sodium buffer for 30 minutes to expose epitopes and enhance antibody binding. The sections were then blocked with an Avidin/Biotin Blocking Kit (Vector Laboratories, SP-2001) and 5% goat serum to prevent the non-specific binding of antibodies to biotin or other proteins.

After blocking, the tissue was incubated with primary anti-F4/80 antibodies at 4°C overnight to allow specific binding to macrophage antigens. The sections were treated with Biotinylated Goat Anti-Rabbit IgG Antibody (Vector Laboratories, BA-1000) at room temperature for 60 minutes, followed by incubation with Streptavidin-Horseradish Peroxidase (Vector Laboratories, SA-5704) for 30 minutes to amplify the staining signal. The staining signal was developed using diaminobenzidine (DAB, Vector Laboratories, SK-4100), producing dark brown staining to mark macrophages. The sections were then counterstained with hematoxylin to visualize cell nuclei, and images were captured using a Nikon Eclipse 80i microscope.

## Supporting information

**S1 Fig: Integrated UMAP plots split by the 5Ht (A) and 6Ho (B) mouse groups.**

**S2 Fig: Subclusters of the 5Ht and 6Ho mice revealed by hierarchical clustering with dendrograms.**

(A) Heatmap showing the dendrograms of the basal epithelial subclusters and their regulon patterns in the 5Ht mice. A branch in the dendrogram showing subcluster S2 is highlighted. (B) Heatmap showing the dendrogram of the basal epithelial subclusters of the 6Ho mice. Cells are shown in rows; the 222 common regulons detected in both mouse groups are shown in columns.

**S3 Fig: Dot plots of the DEGs between subcluster S2 of the 5Ht and 6Ho mice enriched in the HIF-1 signaling pathway.**

Subcluster S2 is highlighted in blue boxes. * indicates DEGs that are either TFs regulating significantly different regulons or their DETGs.

**S4 Fig: Dot plots of the DEGs between subcluster S2 of the 5Ht and 6Ho mice, enriched in various stem cell-related pathways.**

(A) stem cell differentiation; (B) stem cell maintenance. Subcluster S2 is highlighted in blue boxes. * indicates DEGs that are either TFs regulating significantly different regulons or their DETGs.

**S5 Fig: Regulons detected in subcluster S2 of the 5Ht mice with the DETGs enriched in the stem cell development pathway.**

Dot plots of the (A) Egr1 and (B) Junb regulons and their DETGs between subcluster S2 of the 5Ht and 6Ho mice are shown. Subcluster S2 is highlighted in blue boxes. The DETGs are sorted by their ranks within their respective 5Ht regulon.

**S6 Fig: Regulons detected in subcluster S2 of the 6Ho mice with the DETGs enriched in the stem cell development pathway.**

Dot plots show the (A) Egr1, (B) Junb, (C) Rel, (D) Irf1, (E) Elf1, and (F) Maf regulons and their DETGs between subcluster S2 of the 5Ht and 6Ho mice. Subcluster S2 is highlighted in blue boxes. The DETGs are sorted by their ranks within their respective 6Ho regulon.

**S1 Table: Numbers of single cells for three major cell types in the 5Ht and 6Ho mice.**

**S2 Table: Transcription factors (TFs) detected in mammary basal epithelial cells of the (A) 5Ht and (B) 6Ho mice.**

These TFs were validated using the MGI database (see Methods).

**S3 Table: The number of differentially expressed genes (DEGs) and the number of significantly different regulons in each subcluster of basal epithelial cells between the 5Ht and 6Ho mice.**

**S4 Table: DEGs enriched with genes involved in key signaling processes in subcluster U1_wt of the 5Ht mice.**

**S5 Table: DEGs enriched with genes involved in key signaling processes in subcluster S2 of the 5Ht and 6Ho mice.**

**S6 Table: Significantly different regulons and their DETGs enriched in stem cell development and differentiation in subcluster U1_wt of the 5Ht mice.**

**S7 Table: Significantly different regulons and their DETGs enriched in inflammatory and immune responses in subcluster U1_wt of the 5Ht mice.**

**S8 Table: Significantly different regulons and their DETGs enriched in stem cell differentiation in subcluster S2 of the 5Ht and 6Ho mice.**

## References

1. Rappu P, Salo AM, Myllyharju J, Heino J. Role of prolyl hydroxylation in the molecular interactions of collagens. Essays in biochemistry. 2019;63(3):325–35.

2. Xu R. P4HA1 is a new regulator of the HIF-1 pathway in breast cancer. Cell stress. 2019;3(1):27.

3. Xu Y, Xia D, Huang K, Liang M. Hypoxia-induced P4HA1 overexpression promotes post-ischemic angiogenesis by enhancing endothelial glycolysis through downregulating FBP1. J Transl Med. 2024;22(1):74.

4. Li M, Wu F, Zheng Q, Wu Y, Wu Ya. Identification of potential diagnostic and prognostic values of P4HA1 expression in lung cancer, breast cancer, and head and neck cancer. DNA and Cell Biology. 2020;39(5):909–17.

5. Xiong G, Stewart RL, Chen J, Gao T, Scott TL, Samayoa LM, et al. Collagen prolyl 4-hydroxylase 1 is essential for HIF-1α stabilization and TNBC chemoresistance. Nature communications. 2018;9(1):4456.

6. Blencowe M, Arneson D, Ding J, Chen Y-W, Saleem Z, Yang X. Network modeling of single-cell omics data: challenges, opportunities, and progresses. Emerging topics in life sciences. 2019;3(4):379–98.

7. McCalla SG, Fotuhi Siahpirani A, Li J, Pyne S, Stone M, Periyasamy V, et al. Identifying strengths and weaknesses of methods for computational network inference from single-cell RNA-seq data. G3: Genes, Genomes, Genetics. 2023;13(3):jkad004.

8. Zhang R, Ren Z, Chen W. SILGGM: An extensive R package for efficient statistical inference in large-scale gene networks. PLoS computational biology. 2018;14(8):e1006369.

9. Yi H, Zhang Q, Lin C, Ma S. Information-incorporated Gaussian graphical model for gene expression data. Biometrics. 2022;78(2):512–23.

10. Chan TE, Stumpf MP, Babtie AC. Gene regulatory network inference from single-cell data using multivariate information measures. Cell systems. 2017;5(3):251–67. e3.

11. Van Dijk D, Sharma R, Nainys J, Yim K, Kathail P, Carr AJ, et al. Recovering gene interactions from single-cell data using data diffusion. Cell. 2018;174(3):716–29. e27.

12. Lim CY, Wang H, Woodhouse S, Piterman N, Wernisch L, Fisher J, et al. BTR: training asynchronous Boolean models using single-cell expression data. BMC bioinformatics. 2016;17:1–18.

13. Zhang S, Pyne S, Pietrzak S, Halberg S, McCalla SG, Siahpirani AF, et al. Inference of cell type-specific gene regulatory networks on cell lineages from single cell omic datasets. Nature Communications. 2023;14(1):3064.

14. Aibar S, González-Blas CB, Moerman T, Huynh-Thu VA, Imrichova H, Hulselmans G, et al. SCENIC: single-cell regulatory network inference and clustering. Nature methods. 2017;14(11):1083–6.

15. Naik S, Larsen SB, Cowley CJ, Fuchs E. Two to tango: dialog between immunity and stem cells in health and disease. Cell. 2018;175(4):908–20.

16. Wu B, Shi X, Jiang M, Liu H. Cross-talk between cancer stem cells and immune cells: potential therapeutic targets in the tumor immune microenvironment. Molecular Cancer. 2023;22(1):38.

17. Bayik D, Lathia JD. Cancer stem cell–immune cell crosstalk in tumour progression. Nature Reviews Cancer. 2021;21(8):526–36.

18. Sipos F, Műzes G. Cancer stem cell relationship with pro-tumoral inflammatory microenvironment. Biomedicines. 2023;11(1):189.

19. Satija R, Farrell JA, Gennert D, Schier AF, Regev A. Spatial reconstruction of single-cell gene expression data. Nature biotechnology. 2015;33(5):495–502.

20. McInnes L, Healy J, Melville J. Umap: Uniform manifold approximation and projection for dimension reduction. arXiv preprint arXiv:180203426. 2018.

21. Pal B, Chen Y, Milevskiy MJ, Vaillant F, Prokopuk L, Dawson CA, et al. Single cell transcriptome atlas of mouse mammary epithelial cells across development. Breast Cancer Research. 2021;23(1):69.

22. Bach K, Pensa S, Grzelak M, Hadfield J, Adams DJ, Marioni JC, et al. Differentiation dynamics of mammary epithelial cells revealed by single-cell RNA sequencing. Nature communications. 2017;8(1):1–11.

23. Zeisel A, Muñoz-Manchado AB, Codeluppi S, Lönnerberg P, La Manno G, Juréus A, et al. Cell types in the mouse cortex and hippocampus revealed by single-cell RNA-seq. Science. 2015;347(6226):1138-42.

24. Baldarelli RM, Smith CL, Ringwald M, Richardson JE, Bult CJ. Mouse Genome Informatics: an integrated knowledgebase system for the laboratory mouse. Genetics. 2024;227(1):iyae031.

25. Magar AG, Morya VK, Kwak MK, Oh JU, Noh KC. A Molecular Perspective on HIF-1α and Angiogenic Stimulator Networks and Their Role in Solid Tumors: An Update. International Journal of Molecular Sciences. 2024;25(6):3313.

26. Niwa H. The principles that govern transcription factor network functions in stem cells. Development. 2018;145(6):dev157420.

27. Naqvi S, Kim S, Hoskens H, Matthews HS, Spritz RA, Klein OD, et al. Precise modulation of transcription factor levels identifies features underlying dosage sensitivity. Nature genetics. 2023;55(5):841–51.

28. Truong AH, Ben-David Y. The role of Fli-1 in normal cell function and malignant transformation. Oncogene. 2000;19(55):6482–9.

29. Wilson NK, Foster SD, Wang X, Knezevic K, Schütte J, Kaimakis P, et al. Combinatorial transcriptional control in blood stem/progenitor cells: genome-wide analysis of ten major transcriptional regulators. Cell stem cell. 2010;7(4):532–44.

30. Li Y, Luo H, Liu T, Zacksenhaus E, Ben-David Y. The ets transcription factor Fli-1 in development, cancer and disease. Oncogene. 2015;34(16):2022–31.

31. Zeng G, Wang T, Zhang J, Kang YJ, Feng L. FLI1 mediates the selective expression of hypoxia-inducible factor 1 target genes in endothelial cells under hypoxic conditions. FEBS Open Bio. 2021;11(8):2174–85.

32. Sakurai T, Kondoh N, Arai M, Hamada Ji, Yamada T, Kihara-Negishi F, et al. Functional roles of Fli-1, a member of the Ets family of transcription factors, in human breast malignancy. Cancer science. 2007;98(11):1775–84.

33. Carron EC, Homra S, Rosenberg J, Coffelt SB, Kittrell F, Zhang Y, et al. Macrophages promote the progression of premalignant mammary lesions to invasive cancer. Oncotarget. 2017;8(31):50731.

34. Dos Anjos Cassado A. F4/80 as a Major Macrophage Marker: The Case of the Peritoneum and Spleen. Results Probl Cell Differ. 2017;62:161–79.

35. Gilmore TD, Gerondakis S. The c-Rel transcription factor in development and disease. Genes & cancer. 2011;2(7):695–711.

36. Schmitt C, Tonnelle C, Dalloul A, Chabannon C, Debre P, Rebollo A. Aiolos and Ikaros: regulators of lymphocyte development, homeostasis and lymphoproliferation. Apoptosis. 2002;7:277–84.

37. Deng Y, Lu L, Zhang H, Fu Y, Liu T, Chen Y. The role and regulation of Maf proteins in cancer. Biomarker Research. 2023;11(1):17.

38. Fan Y, Mao R, Yang J. NF-κB and STAT3 signaling pathways collaboratively link inflammation to cancer. Protein & cell. 2013;4:176–85.

39. Hoesel B, Schmid JA. The complexity of NF-κB signaling in inflammation and cancer. Molecular cancer. 2013;12:1–15.

40. Islam Z, Ali AM, Naik A, Eldaw M, Decock J, Kolatkar PR. Transcription factors: The fulcrum between cell development and carcinogenesis. Frontiers in oncology. 2021;11:681377.

41. Liang Q, Xu C, Chen X, Li X, Lu C, Zhou P, et al. The roles of Mesp family proteins: functional diversity and redundancy in differentiation of pluripotent stem cells and mammalian mesodermal development. Protein & cell. 2015;6(8):553–61.

42. Wolf FA, Angerer P, Theis FJ. SCANPY: large-scale single-cell gene expression data analysis. Genome biology. 2018;19:1–5.

43. Moerman T, Aibar Santos S, Bravo González-Blas C, Simm J, Moreau Y, Aerts J, et al. GRNBoost2 and Arboreto: efficient and scalable inference of gene regulatory networks. Bioinformatics. 2019;35(12):2159–61.

44. Finak G, McDavid A, Yajima M, Deng J, Gersuk V, Shalek AK, et al. MAST: a flexible statistical framework for assessing transcriptional changes and characterizing heterogeneity in single-cell RNA sequencing data. Genome biology. 2015;16:1–13.

45. Benjamini Y, Hochberg Y. Controlling the false discovery rate: a practical and powerful approach to multiple testing. Journal of the Royal statistical society: series B (Methodological). 1995;57(1):289–300.

46. Wu T, Hu E, Xu S, Chen M, Guo P, Dai Z, et al. clusterProfiler 4.0: A universal enrichment tool for interpreting omics data. Innovation (Camb). 2021;2(3):100141.

